# The relationship between passive ankle joint stiffness and the stiffness of muscles, nerve, and tendon

**DOI:** 10.1101/2025.09.09.675058

**Authors:** Hiyu Mukai, Jun Umehara, Junya Saeki, Ko Yanase, Zimin Wang, Hiroshige Tateuchi, Noriaki Ichihashi

**Affiliations:** Human Health Sciences, Graduate School of Medicine, Kyoto University, 53 Shogoin-Kawahara-cho, Kyoto 606-8507, Japan; Faculty of Rehabilitation, Kansai Medical University, 18-89 Uyama-higashi, Hirakata, Osaka 573-1136, Japan; Graduate School of Rehabilitation, Osaka Kawasaki Rehabilitation University, 158 Mizuma, Kaizuka, Osaka 597-0104, Japan; Faculty of Health and Sports Science, Doshisha University, 1-3 Tatara-Miyakodani, Kyotanabe, Kyoto 610-0394, Japan

**Keywords:** Joint flexibility, stiffness, triceps surae muscles, tibial nerve, Achilles tendon

## Abstract

Passive joint stiffness reflects the stiffness of various soft tissues across a joint. However, no previous studies have investigated the relationship between passive joint stiffness and muscle, nerve, and tendon stiffness. This study aimed to clarify whether passive ankle joint stiffness is related to stiffness in the triceps surae muscles, tibial nerve, and Achilles tendon. Thirty-eight healthy adults participated in the study. The passive ankle joint stiffness (slope of angle–passive torque curve) and shear wave velocities, which indicate soft tissue stiffness, of the triceps surae muscles and tibial nerve were measured at 5° of ankle plantarflexion and 5°, 15°, and 25° of ankle dorsiflexion. The shear wave velocity of the Achilles tendon was measured only at 5° of plantarflexion. A multiple regression model (forced-entry method) was constructed at each angle, specifying the shear wave velocities as the independent variables and passive joint stiffness as the dependent variable. At 5° of plantarflexion, no shear wave velocities were significantly related to passive joint stiffness (all *p ≥* 0.05). At 5° and 15° of dorsiflexion, only the shear wave velocities of the tibial nerve were significantly positively related to passive joint stiffness (*p* = 0.024 and 0.008, respectively). At 25° of dorsiflexion, the shear wave velocities of the lateral gastrocnemius muscle and tibial nerve were significantly positively related to passive joint stiffness (*p* = 0.002 and 0.001, respectively). It can be concluded that both triceps surae muscles stiffness and tibial nerve stiffness are related to passive ankle joint stiffness.

## 1. Introduction

Joint flexibility is regarded as an important outcome in sports and rehabilitation settings. The maximum angle during passive movement is often used as an indicator of flexibility. However, this measure reflects not only the flexibility of the soft tissues surrounding the joint but also stretch tolerance (Ingram et al., 2025; Weppler and Magnusson, 2010). Therefore, researchers have adopted passive joint stiffness, which is the slope of the angle–passive joint torque curve, as an indicator of joint flexibility (Chesworth and Vandervoort, 1989; Riemann et al., 2001). Previous studies have reported that an increase in passive ankle joint stiffness was observed in patients with spastic hypertonia (Chung et al., 2004) and diabetes mellitus (Rao et al., 2006) and in patients after ankle cast removal (Chesworth and Vandervoort, 1995). An increase in passive joint stiffness may lead to activity limitations such as difficulty walking and it is important to investigate the factors relevant to passive joint stiffness to develop effective approaches for reducing stiffness.

Passive joint stiffness is considered to reflect the stiffness of various soft tissues across a joint (Riemann et al., 2001; Takeuchi et al., 2023). Some previous studies have investigated the relationship between passive joint stiffness and muscle and tendon stiffness using shear wave elastography. The results indicate that in healthy young adults, passive joint stiffness is not related to medial gastrocnemius and soleus muscle stiffness at 0° of ankle dorsiflexion; however, it is significantly positively related to triceps surae muscles stiffness at 15° or 20° of ankle dorsiflexion (Chino and Takahashi, 2016, 2015; Hirata and Akagi, 2022). Chino and Takahashi (2015) noted no significant correlation between passive joint stiffness and Achilles tendon stiffness at 0° of ankle dorsiflexion. Although there may be correlations between the stiffness of each soft tissue, previous studies have not investigated whether the stiffness of each soft tissue is independently related to passive joint stiffness. Additionally, in recent studies, researchers have evaluated nerve stiffness by using shear wave elastography and reported a significant relationship between tibial nerve stiffness and the maximum angle of ankle dorsiflexion (Kawanishi et al., 2022; Mukai et al., 2025, preprint). However, the exact relationship between passive joint stiffness and tibial nerve stiffness remains unclear. Therefore, it is necessary to investigate whether stiffness in the triceps surae muscles, Achilles tendon, and tibial nerve are independently related to passive joint stiffness, considering the correlation among soft tissue stiffnesses.

This study aimed to clarify whether triceps surae (medial and lateral gastrocnemius and soleus) muscles, Achilles tendon, and tibial nerve stiffness are independently related to passive ankle joint stiffness using shear wave elastography. Passive joint stiffness is considered to reflect the stiffness of various soft tissues around a joint (Riemann et al., 2001; Takeuchi et al., 2023). Previous studies have reported no significant correlation between passive joint stiffness and the medial gastrocnemius and soleus muscles at slightly lengthened positions. In contrast, significant correlations have been observed at greatly lengthened positions (Chino and Takahashi, 2016; Hirata and Akagi, 2022). We hypothesized that i) passive ankle joint stiffness is related to triceps surae muscles, Achilles tendon, and tibial nerve stiffness, and that ii) triceps surae muscles stiffness is related to passive ankle joint stiffness more strongly at greatly lengthened positions.

## 2. Methods

### 2.1. Participants

Forty-three healthy adults with no pain or limitation in the range of motion in their ankle joint on the non-dominant side participated in this study (19 males and 24 females; age, 24.3 ± 2.9 years; height, 165.8 ± 7.7 cm; body mass, 59.3 ± 8.2 kg). When the non-dominant leg was defined as the leg opposite to the one used in kicking a ball, 4 right legs and 39 left legs were targeted. All participants were informed regarding the study procedure in advance and written consent for participation was obtained. This study complied with the Declaration of Helsinki and was approved by the Ethics Committee (approval day: March 26, 2024, approval number: C1652-2).

### 2.2. Experimental procedure

This cross-sectional study investigated the relationship between passive ankle joint stiffness and the shear wave velocities of the medial and lateral gastrocnemius muscles, soleus muscle, tibial nerve, and Achilles tendon, which were considered to represent the stiffness of these soft tissues. To measure these values, the participants were laid in a prone position on the seat of dynamometer (BIODEX System 4, BIODEX, NY, USA), with the hip and knee joint in neutral positions, foot of non-dominant side fixed to the footplate, and foot and pelvis fixed by straps. Participants were asked to relax during measurement. The measurement of the shear wave velocities was followed by that of the passive joint torque, which was necessary to calculate the passive joint stiffness.

### 2.3. Measurement of shear wave velocities

The shear wave velocities were measured using the shear wave elastography mode in an ultrasound system (Aixplorer v12.2, SuperSonic Imagine, Aix-en-Provence, France). A linear probe (2 to 10 MHz, SuperLinear SL10-2) and musculoskeletal preset (muscle mode) were used to measure the shear wave velocities of the triceps surae muscles and tibial nerve. Another linear probe (4 to 15 MHz, SuperLinear SL 15-4) and musculoskeletal preset (foot–ankle mode) were used to measure the shear wave velocity of the Achilles tendon. All measurements were performed with the following settings: mode, penetration; frequency, 1.7 Hz; smoothing level, five; persistence, high; opacity, 100%. We interpreted higher shear wave velocities as representing higher stiffness. The shear wave velocities were measured at the following regions: the medial and lateral gastrocnemius muscles at 30% of lower leg length (Akagi and Takahashi, 2013; Nakamura et al., 2014; Taniguchi et al., 2015), soleus muscle at 50% of lower leg length (Kubo et al., 2017), tibial nerve at the region near the medial malleolus (determined through a preliminary study), and Achilles tendon at 4 cm proximal to the insertion to the calcaneal tubercle (Selcuk Can et al., 2022). To minimize the effect of stretching, the shear wave velocities were measured at 5° of ankle plantarflexion and 5°, 15°, and 25° of ankle dorsiflexion in order. The shear wave velocity of the Achilles tendon was measured only at 5° of ankle plantarflexion due to saturation.

To analyze the ultrasound images, a rectangular region of interest 1 cm in height and 2 cm in width was defined. Then, using a Q-box trace function, the maximum area within the region of interest was encapsulated, excluding the bone, aponeurosis, or epineurium, and the mean of the shear wave velocities in the encapsulated region was calculated. Two images of each muscle and three images of the tibial nerve and Achilles tendon were acquired, and the mean values were used in our statistical analysis. The analyzed ultrasound images are presented in Figure 1.

**Figure 1:**
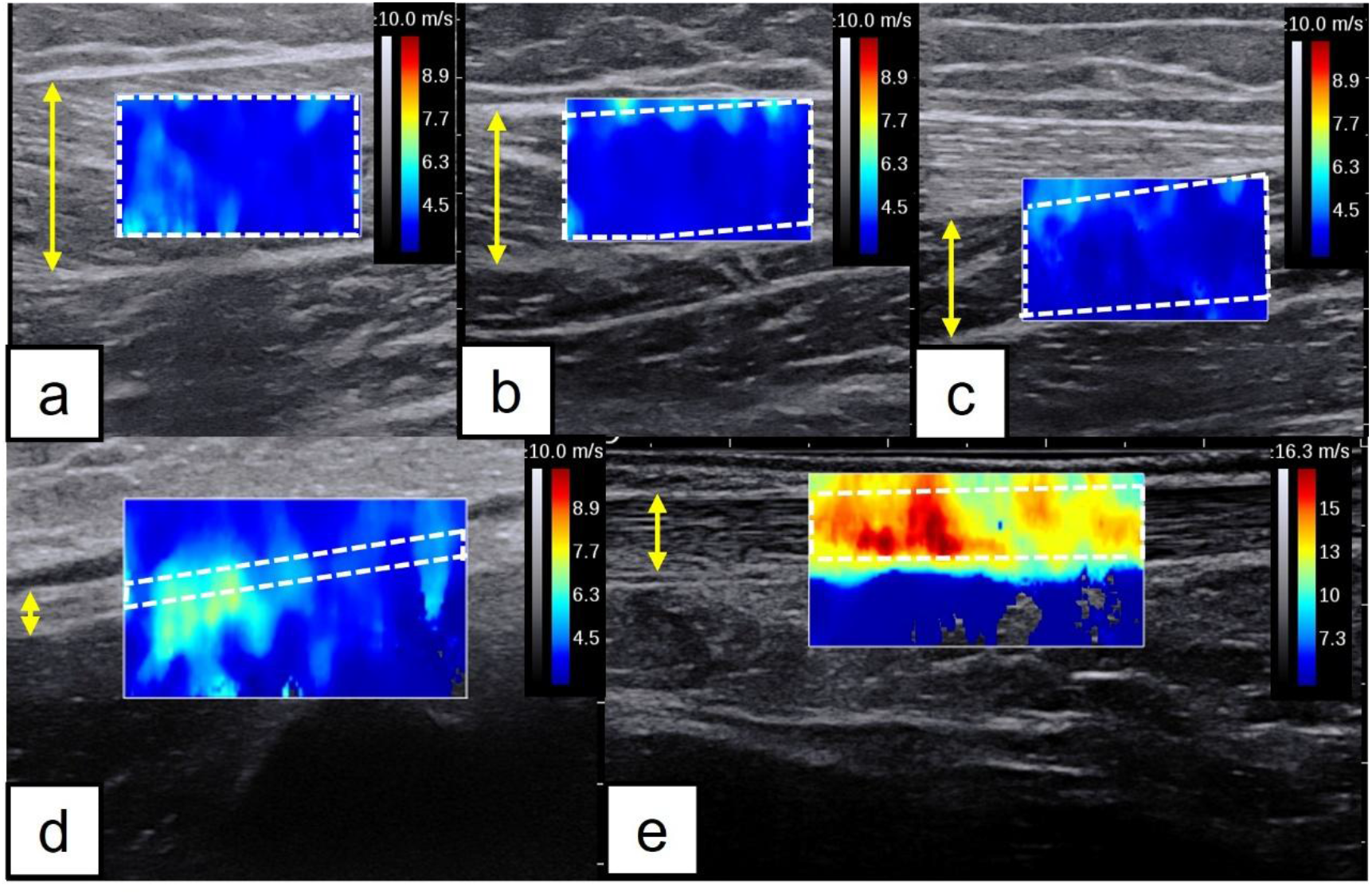
Analyzed ultrasound images a: medial gastrocnemius muscle, b: lateral gastrocnemius muscle, c: soleus muscle, d: tibial nerve, and e: Achilles tendon. The rectangle in the ultrasound image is the region of interest. The area surrounded by white wavy lines is the target area of analysis. The area indicated by the yellow arrow is the target tissue for measurement.

### 2.4. Measurement and correction of passive joint torque and calculation of passive joint stiffness

To measure the passive joint torque, each participant’s ankle was passively dorsiflexed by 5°/s from 30° of ankle plantarflexion to the maximum dorsiflexion angle, where the participant felt discomfort without pain (Nakamura et al., 2017). Prior to the testing session, two sessions were conducted to familiarize the participants with the procedure (Hirata et al., 2015; Konrad et al., 2015; Konrad and Tilp, 2014). Then, three testing sessions were conducted. To subtract the torque generated by the mass of foot from the measured passive joint torque, the gravitational force acting on the foot was calculated by dividing the measured passive joint torque at 30° of ankle plantarflexion by sine(30°). Because 30° of ankle plantarflexion is under slack angle for the triceps surae muscles (Hirata et al., 2016), the measured passive joint torque at this angle can be interpreted as the torque generated by the mass of foot. The torque generated by the mass of foot at any angle of ankle plantarflexion/dorsiflexion was then calculated by multiplying the gravitational force acting on the foot by the sine of the angle. The passive joint torque was corrected by subtracting the torque generated by the mass of foot from the measured passive joint torque. The relationship between the angle and corrected passive joint torque was generated by fitting a fourth-order polynomial equation to the data (*y = ax*^*4*^ *+ bx*^*3*^ *+ cx*^*2*^ *+ dx + e*, where *y* is the torque, *x* is the joint angle, and *a* to *e* are constants). The passive joint stiffness at 5° of ankle plantarflexion, and 5°, 15°, and 25° of ankle dorsiflexion was calculated using the first derivative (slope) of that equation (Chesworth and Vandervoort, 1989; Riemann et al., 2001). The passive joint stiffness was calculated three times and the mean value was used for statistical analysis.

### 2.5. Statistical analysis

SPSS Statistics 22 (IBM, Armonk, NY, USA) was used for statistical analysis. To confirm the normality of the data, the Shapiro–Wilk test was performed. As a primary analysis, to clarify the relationship between the soft tissue stiffness and passive joint stiffness, multiple regression analysis using the forced-entry method was performed, specifying the shear wave velocity of each soft tissue as the independent variable and passive joint stiffness as the dependent variable. A regression model was constructed at each angle for which the shear wave velocities and passive joint stiffness were calculated. As a sub-analysis, to clarify the correlation in the shear wave velocities among the soft tissues at 5°, 15°, and 25° of ankle dorsiflexion, Pearson’s product–moment correlation coefficients or Spearman’s rank correlation coefficients were calculated. Statistical significance for all results was defined as *p* < 0.05.

## 3. Results

### 3.1. Characteristics of participants

After excluding four participants whose maximum ankle dorsiflexion angle was less than 25° and one participant who felt pain during the measurement of the shear wave velocities, 38 participants were included in the statistical analysis. The characteristics of the 38 participants included in the statistical analysis were as follows: 17 males and 21 females; age, 24.6 ± 2.8 years; height, 166.2 ± 7.6 cm; body mass, 60.1 ± 8.0 kg; 33 right-footed and 5 left-footed individuals. The passive joint stiffness and shear wave velocities at each angle are presented in Table 1.

**Table 1:** Passive ankle joint stiffness and shear wave velocity of each tissue.

|  |  | 5° of ankle<br>plantarflexion | 5° of ankle<br>dorsiflexion | 15° of ankle<br>dorsiflexion | 25° of ankle<br>dorsiflexion |
| --- | --- | --- | --- | --- | --- |
| Passive ankle joint stiffness (Nm/°) |  | 0.005 ± 0.075 | 0.182 ± 0.147 | 0.506 ± 0.244 | 0.969 ± 0.456 |
| Shear wave<br>velocity (m/s) | Medial<br>gastrocnemius | 3.362 ± 0.440 | 4.607 ± 0.637 | 6.486 ± 0.802 | 8.379 ± 0.840 |
|  | Lateral<br>gastrocnemius | 3.300 ± 0.576 | 4.041 ± 0.689 | 5.234 ± 0.697 | 6.914 ± 0.921 |
|  | Soleus | 3.139 ± 0.823 | 3.271 ± 0.864 | 3.459 ± 0.777 | 3.982 ± 0.970 |
|  | Tibial nerve | 5.782 ± 1.284 | 6.556 ± 1.295 | 7.055 ± 1.361 | 7.632 ± 1.219 |
|  | Achilles tendon | 14.839 ± 0.650 | N/A | N/A | N/A |

Variables are described as mean ± SD.

### 3.2. Relationship between passive joint stiffness and shear wave velocity

At 5° of ankle plantarflexion, no shear wave velocities of any soft tissues were significantly related to passive joint stiffness (all *p* ≥ 0.05). At 5° and 15° of ankle dorsiflexion, the shear wave velocities of the tibial nerve were significantly positively related to passive joint stiffness (*p* = 0.024 and 0.008, respectively). However, the shear wave velocities of the triceps surae muscles were not significantly related to passive joint stiffness (all *p* ≥ 0.05). At 25° of ankle dorsiflexion, the shear wave velocities of the lateral gastrocnemius muscle and tibial nerve were significantly positively related to passive joint stiffness (*p* = 0.002 and 0.001, respectively). The shear wave velocities of the medial gastrocnemius and soleus muscles were not significantly related to passive joint stiffness (*p* = 0.185 and 0.394, respectively). The results of multiple regression analysis at each angle are presented in Table 2.

**Table 2:** Results of multiple regression analysis.

| 5° of ankle plantarflexion (adjusted R<sup>2</sup> = 0.152) |  |  |  |  |  |
| --- | --- | --- | --- | --- | --- |
|  | B | β | 95%CI | *p* value | VIF |
| Medial gastrocnemius | −0.044 | −0.256 | −0.113 to 0.025 | 0.204 | 1.704 |
| Lateral gastrocnemius | 0.051 | 0.388 | −0.003 to 0.105 | 0.064 | 1.785 |
| Soleus | < 0.001 | 0.004 | −0.032 to 0.033 | 0.980 | 1.308 |
| Tibial nerve | 0.015 | 0.259 | −0.006 to 0.036 | 0.148 | 1.334 |
| Achilles tendon | 0.021 | 0.180 | −0.019 to 0.061 | 0.295 | 1.249 |
| 5° of ankle dorsiflexion (adjusted R<sup>2</sup> = 0.101) |  |  |  |  |  |
|  | B | β | 95%CI | *p* value | VIF |
| Medial gastrocnemius | −0.070 | −0.304 | −0.157 to 0.016 | 0.109 | 1.402 |
| Lateral gastrocnemius | 0.044 | 0.206 | −0.040 to 0.127 | 0.294 | 1.535 |
| Soleus | −0.006 | −0.037 | −0.067 to 0.054 | 0.833 | 1.255 |
| Tibial nerve | 0.044 | 0.385 | 0.006 to 0.081 | 0.024* | 1.086 |
| 15° of ankle dorsiflexion (adjusted R<sup>2</sup> = 0.198) |  |  |  |  |  |
|  | B | β | 95%CI | *p* value | VIF |
| Medial gastrocnemius | 0.040 | 0.131 | −0.091 to 0.171 | 0.540 | 2.057 |
| Lateral gastrocnemius | 0.024 | 0.070 | −0.143 to 0.192 | 0.769 | 2.541 |
| Soleus | −0.040 | −0.126 | −0.143 to 0.064 | 0.443 | 1.205 |
| Tibial nerve | 0.088 | 0.489 | 0.025 to 0.151 | 0.008* | 1.377 |
| 25° of ankle dorsiflexion (adjusted R<sup>2</sup> = 0.411) |  |  |  |  |  |
|  | B | β | 95%CI | *p* value | VIF |
| Medial gastrocnemius | −0.103 | −0.190 | −0.258 to 0.052 | 0.185 | 1.238 |
| Lateral gastrocnemius | 0.240 | 0.484 | 0.095 to 0.384 | 0.002* | 1.286 |
| Soleus | −0.053 | −0.113 | −0.178 to 0.072 | 0.394 | 1.077 |
| Tibial nerve | 0.176 | 0.470 | 0.076 to 0.275 | 0.001* | 1.072 |

B: unstandardized regression coefficient, β: standardized regression coefficient, CI: confidence
interval, VIF: variance inflation factor, \*: *p* < 0.05.

### 3.3. Correlation of shear wave velocities between each soft tissue

At 5° of ankle dorsiflexion, the shear wave velocity of the lateral gastrocnemius muscle was significantly correlated with those of the medial gastrocnemius (ρ = 0.591, *p* < 0.001) and soleus muscles (ρ = 0.501, *p* = 0.001). At 15° of ankle dorsiflexion, the shear wave velocity of the medial gastrocnemius muscle was significantly correlated with those of the lateral gastrocnemius (r = 0.666, *p* < 0.001) and soleus muscles (ρ = 0.384, *p* = 0.017). Additionally, the shear wave velocity of the lateral gastrocnemius muscle was significantly correlated with those of the soleus muscle (ρ = 0.465, *p* = 0.003) and tibial nerve (r = 0.423, *p* = 0.008). At 25° of ankle dorsiflexion, the shear wave velocity of the medial gastrocnemius muscle was significantly correlated with that of the lateral gastrocnemius muscle (r = 0.426, *p* = 0.008). All results for the correlation of the shear wave velocities between each soft tissue are provided in Supplemental Material 1.

## 4. Discussion

In this study, we investigated the relationship between passive joint stiffness and the shear wave velocities of the triceps surae muscles, Achilles tendon, and tibial nerve. We found that the shear wave velocity of the tibial nerve was independently related to passive joint stiffness at 5°, 15°, and 25° of ankle dorsiflexion. At 25° of ankle dorsiflexion, the shear wave velocity of the lateral gastrocnemius muscle is also independently related to passive joint stiffness. Previous studies have reported significant relationships between maximum joint angle (an indicator of joint flexibility), and muscle (Hirata et al., 2020; Miyamoto et al., 2018) and nerve stiffness (Kawanishi et al., 2022; Mukai et al., 2025, preprint). This is the first study to demonstrate that muscle and nerve stiffness are also related to passive joint stiffness.

The positive relationship between passive joint stiffness and the shear wave velocity of the tibial nerve at 5°, 15°, and 25° of ankle dorsiflexion indicates that individuals with higher tibial nerve stiffness have higher joint stiffness. Although passive joint stiffness has been suggested to reflect the stiffness of not only muscles but also other soft tissues (Riemann et al., 2001; Takeuchi et al., 2023), this assumption has not been clearly validated through empirical studies. It should be noted that a significant relationship with tibial nerve stiffness was observed at 5° and 15° of ankle dorsiflexion; however, the adjusted R^2^ values were small (0.101 and 0.198, respectively). Therefore, passive joint stiffness cannot be explained by tibial nerve stiffness alone.

Contradictory to our two hypotheses, excluding the lateral gastrocnemius muscle at 25° of ankle dorsiflexion, the shear wave velocities of the triceps surae muscles were not significantly related to passive joint stiffness. Chino and Takahashi (2015) reported no significant relationship between the shear modulus of the medial gastrocnemius muscle and passive joint stiffness at a slightly lengthened position, which is consistent with the results of this study. However, inconsistent with this study, previous studies have reported that the shear moduli of the triceps surae muscles were significantly positively related to passive joint stiffness (Chino and Takahashi, 2016; Hirata and Akagi, 2022). Koo and Hug (2015) proposed the following equation:

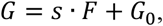

where *G* is the muscle shear modulus at any length, *F* is the muscle passive force at the length along its long axis, *G*_*0*_ is the muscle shear modulus at its slack length, and *s* is the slope of the *F–G* relationship. In this study, individual differences may have existed in *G*_*0*_ and the relative proportion of each muscle comprising the triceps surae muscles in *F*. Considering such individual differences, triceps surae muscles stiffness may not be related to passive joint stiffness, excluding the lateral gastrocnemius muscle at 25° of ankle dorsiflexion.

As a result of multiple regression analysis, no significant relationship was observed between passive joint stiffness and the shear wave velocity of the lateral gastrocnemius muscle at 15° of ankle dorsiflexion (Table 2). However, correlation analysis revealed that passive joint stiffness tended to be positively correlated with the shear wave velocity of the lateral gastrocnemius muscle at 15° of ankle dorsiflexion (r = 0.314, *p* = 0.055) (Supplemental Material 2). At this angle, a significant correlation was observed between the shear wave velocities of the lateral gastrocnemius muscle and that of the tibial nerve (r = 0.423, *p* = 0.008) (Supplemental Material 1). These results suggest the necessity for considering the correlations among the stiffness of each soft tissue to investigate the relationship between passive joint stiffness and the stiffness of various soft tissues. Hirata and Akagi (2022) investigated the relationship between triceps surae (medial and lateral gastrocnemius and soleus) muscles stiffness and passive ankle joint stiffness. However, they did not perform statistical analysis considering the correlations among the stiffness values of each muscle. Although the physiological or biomechanical mechanisms associated with the correlation among the stiffness values of each soft tissue remain unclear, the consideration of correlation may provide more meaningful knowledge when investigating the relationship between passive joint stiffness and the stiffness of various soft tissues.

At 5° of ankle plantarflexion, no significant relationship was observed between passive joint stiffness and Achilles tendon stiffness. However, the passive joint stiffness at this angle was minimal (Table 1) and the Achilles tendon may not have been sufficiently elongated. In this study, we could not measure the shear wave velocity of the Achilles tendon in ankle dorsiflexion due to saturation. Therefore, we cannot say definitively that Achilles tendon stiffness is not related to passive ankle joint stiffness.

There are two limitations of this study. First, we did not evaluate the stiffness of soft tissues other than the triceps surae muscles and tibial nerve, such as other plantarflexors, aponeurosis, skin, blood vessels, or subcutaneous fat. Because passive joint stiffness reflects the stiffness of various soft tissues across the joint, soft tissues which were not evaluated in this study may be related to passive joint stiffness. Second, we evaluated only at the position with 0° of hip and knee flexion. Previous studies have reported that the shear wave velocities of the triceps surae muscles and tibial nerve varied depending on the angles of hip or knee flexion, even at the same angle of ankle dorsiflexion (Andrade et al., 2022; Ateş et al., 2018; Le Sant et al., 2017). For hip or knee flexion angles different from that considered in this study, different results may be observed in the relationship between passive joint stiffness and soft tissues stiffness. In future studies, it will be necessary to investigate these relationships by considering additional soft tissues and positions.

## 5. Conclusion

We investigated the relationship between passive ankle joint stiffness and soft tissue (triceps surae muscles, tibial nerve, and Achilles tendon) stiffness using shear wave elastography. At 5° and 15° of ankle dorsiflexion, only the shear wave velocities of the tibial nerve were significantly related to passive joint stiffness. At 25° of ankle dorsiflexion, the shear wave velocities of the lateral gastrocnemius muscle and tibial nerve were significantly related to passive joint stiffness. This results of this study suggest that passive joint stiffness reflects the stiffness of not only muscle but also nerves.

## CRediT authorship contribution statement

**Hiyu Mukai:** Writing – original draft, Conceptualization, Data curation, Formal analysis, Investigation, Methodology, Visualization. **Jun Umehara:** Conceptualization, Methodology, Writing – review, Editing. **Junya Saeki:** Conceptualization, Methodology, Writing – review, Editing. **Ko Yanase:** Conceptualization, Methodology, Writing – review, Editing. **Zimin Wang:** Conceptualization, Methodology, Writing – review, Editing. **Hiroshige Tateuchi:** Conceptualization, Methodology, Writing – review, Editing. **Noriaki Ichihashi:** Conceptualization, Methodology, Writing – review, Editing. Project administration and Supervision.

## Data statement

The datasets generated during and/or analyzed during this current study are available from the corresponding author upon reasonable request.

## Acknowledgement

We are grateful to Editage (www.editage.jp) for their assistance with English language editing.

---

**Supplemental Material 1:**
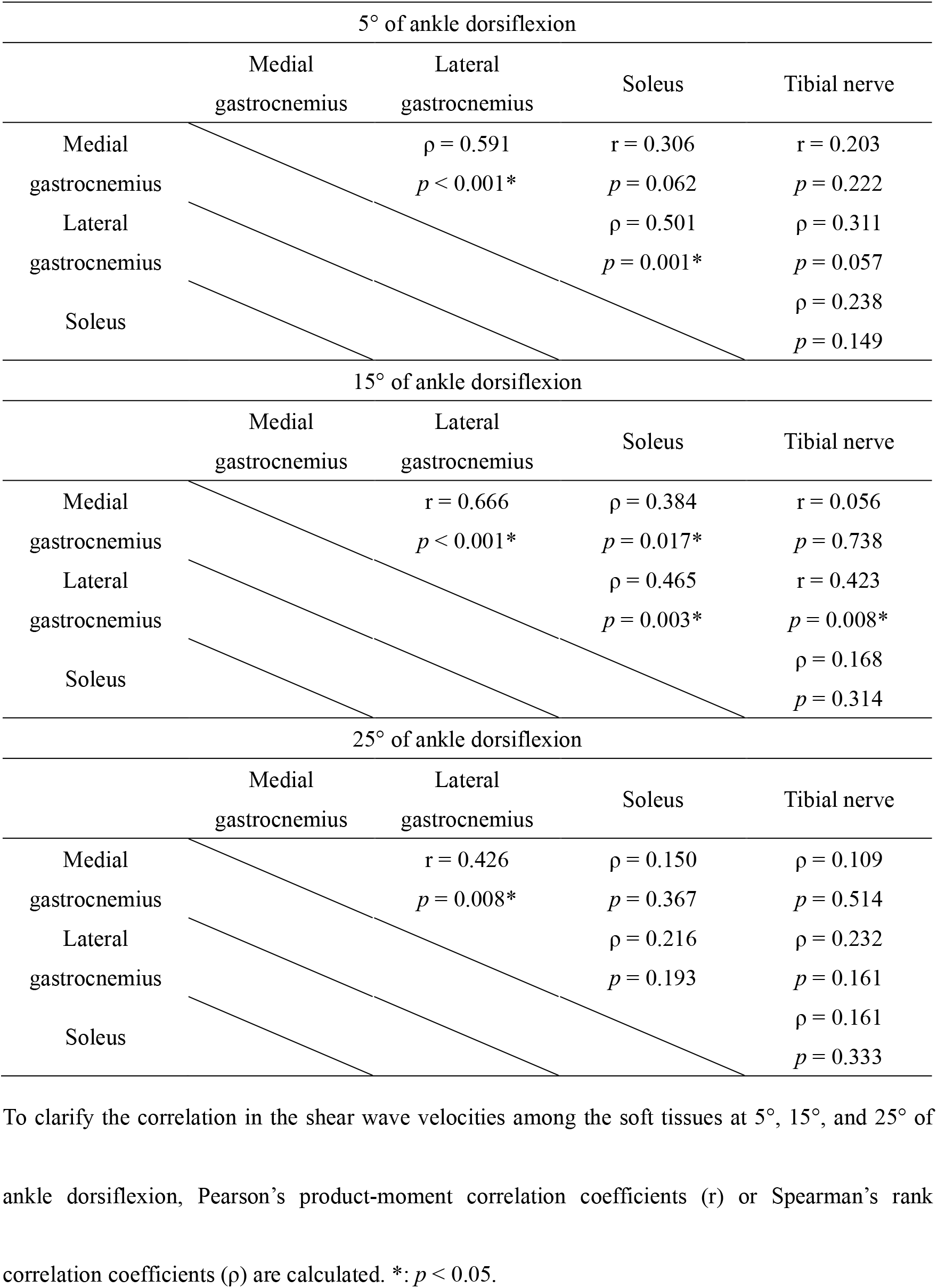
Correlation of shear wave velocity between each tissue.

| 5° of ankle dorsiflexion |  |  |  |  |
| --- | --- | --- | --- | --- |
|  | Medial<br>gastrocnemius | Lateral<br>gastrocnemius | Soleus | Tibial nerve |
| Medial<br>gastrocnemius | | $\rho = 0.591$<br>$p < 0.001^*$ | $r = 0.306$<br>$p = 0.062$ | $r = 0.203$<br>$p = 0.222$ |
| Lateral<br>gastrocnemius | | | $\rho = 0.501$<br>$p = 0.001^*$ | $\rho = 0.311$<br>$p = 0.057$ |
| Soleus | | | | $\rho = 0.238$<br>$p = 0.149$ |
| 15° of ankle dorsiflexion |  |  |  |  |
|  | Medial<br>gastrocnemius | Lateral<br>gastrocnemius | Soleus | Tibial nerve |
| Medial<br>gastrocnemius | | $r = 0.666$<br>$p < 0.001^*$ | $\rho = 0.384$<br>$p = 0.017^*$ | $r = 0.056$<br>$p = 0.738$ |
| Lateral<br>gastrocnemius | | | $\rho = 0.465$<br>$p = 0.003^*$ | $r = 0.423$<br>$p = 0.008^*$ |
| Soleus | | | | $\rho = 0.168$<br>$p = 0.314$ |
| 25° of ankle dorsiflexion |  |  |  |  |
|  | Medial<br>gastrocnemius | Lateral<br>gastrocnemius | Soleus | Tibial nerve |
| Medial<br>gastrocnemius | | $r = 0.426$<br>$p = 0.008^*$ | $\rho = 0.150$<br>$p = 0.367$ | $\rho = 0.109$<br>$p = 0.514$ |
| Lateral<br>gastrocnemius | | | $\rho = 0.216$<br>$p = 0.193$ | $\rho = 0.232$<br>$p = 0.161$ |
| Soleus | | | | $\rho = 0.161$<br>$p = 0.333$ |

To clarify the correlation in the shear wave velocities among the soft tissues at 5°, 15°, and 25° of ankle dorsiflexion, Pearson's product-moment correlation coefficients (r) or Spearman's rank correlation coefficients ( $\rho$ ) are calculated. \*: $p < 0.05$ .

**Supplemental material 2:** Correlation between passive joint stiffness and shear wave velocities at 5°, 15°, and 25° of ankle dorsiflexion.

|  |  | Passive ankle joint stiffness |  |  |
| --- | --- | --- | --- | --- |
|  |  | 5° of ankle dorsiflexion | 15° of ankle dorsiflexion | 25° of ankle dorsiflexion |
| Shear wave velocity | Medial gastrocnemius | $r = -0.126$ | $r = 0.162$ | $\rho = 0.029$ |
| | | $p = 0.451$ | $p = 0.331$ | $p = 0.861$ |
| | Lateral gastrocnemius | $\rho = 0.097$ | $r = 0.314$ | $\rho = 0.479$ |
| | | $p = 0.564$ | $p = 0.055$ | $p = 0.002$ |
| | Soleus | $\rho = 0.229$ | $\rho = 0.041$ | $\rho = 0.034$ |
| | | $p = 0.167$ | $p = 0.805$ | $p = 0.838$ |
| | Tibial nerve | $r = 0.342$ | $r = 0.490$ | $\rho = 0.537$ |
| | | $p = 0.036^*$ | $p = 0.002^*$ | $p = 0.001^*$ |

To clarify the correlation between the shear wave velocity and passive joint stiffness at 5°, 15°, and 25° of ankle dorsiflexion, Pearson's product-moment correlation coefficients ( $r$ ) or Spearman's rank correlation coefficients ( $\rho$ ) are calculated. \*: $p < 0.05$ .

## References

Akagi, R., Takahashi, H., 2013. Acute effect of static stretching on hardness of the gastrocnemius muscle. Med Sci Sports Exerc 45, 1348–1354. 10.1249/MSS.0b013e3182850e17

Andrade, R.J., Freitas, S.R., Hug, F., Coppieters, M.W., Sierra-Silvestre, E., Nordez, A., 2022. Spatial variation in mechanical properties along the sciatic and tibial nerves: An ultrasound shear wave elastography study. J Biomech 136. 10.1016/j.jbiomech.2022.111075

Ateş, F., Andrade, R.J., Freitas, S.R., Hug, F., Lacourpaille, L., Gross, R., Yucesoy, C.A., Nordez, A., 2018. Passive stiffness of monoarticular lower leg muscles is influenced by knee joint angle. Eur J Appl Physiol 118, 585–593. 10.1007/s00421-018-3798-y

Chesworth, B.B., Vandervoort, A.A., 1995. Comparison of Passive Stiffness Variables and Range of Motion in Uninvolved and Involved Ankle Joints of Patients Following Ankle Fractures. Phys Ther 75, 253–261. 10.1093/ptj/75.4.253

Chesworth, B.M., Vandervoort, A.A., 1989. Age and Passive Ankle Stiffiiess in Healthy Women. Phys Ther 69, 217–224. 10.1093/ptj/69.3.217

Chino, K., Takahashi, H., 2016. Measurement of gastrocnemius muscle elasticity by shear wave elastography: association with passive ankle joint stiffness and sex differences. Eur J Appl Physiol 116, 823–830. 10.1007/s00421-016-3339-5

Chino, K., Takahashi, H., 2015. The association of muscle and tendon elasticity with passive joint stiffness: In vivo measurements using ultrasound shear wave elastography. Clinical Biomechanics 30, 1230–1235. 10.1016/j.clinbiomech.2015.07.014

Chung, S.G., Van Rey, E., Bai, Z., Roth, E.J., Zhang, L.Q., 2004. Biomechanic changes in passive properties of hemiplegic ankles with spastic hypertonia. Arch Phys Med Rehabil 85, 1638–1646. 10.1016/j.apmr.2003.11.041

Hirata, K., Akagi, R., 2022. Contribution of muscle stiffness of the triceps surae to passive ankle joint stiffness in young and older adults. Front Physiol 13. 10.3389/fphys.2022.972755

Hirata, K., Kanehisa, H., Miyamoto-Mikami, E., Miyamoto, N., 2015. Evidence for intermuscle difference in slack angle in human triceps surae. J Biomech 48, 1210–1213. 10.1016/j.jbiomech.2015.01.039

Hirata, K., Miyamoto-Mikami, E., Kanehisa, H., Miyamoto, N., 2016. Muscle-specific acute changes in passive stiffness of human triceps surae after stretching. Eur J Appl Physiol 116, 911–918. 10.1007/s00421-016-3349-3

Hirata, K., Yamadera, R., Akagi, R., 2020. Associations between Range of Motion and Tissue Stiffness in Young and Older People. Med Sci Sports Exerc 52, 2179–2188. 10.1249/MSS.0000000000002360

Ingram, L.A., Tomkinson, G.R., d’Unienville, N.M.A., Gower, B., Gleadhill, S., Boyle, T., Bennett, H., 2025. Mechanisms Underlying Range of Motion Improvements Following Acute and Chronic Static Stretching: A Systematic Review, Meta-analysis and Multivariate Metaregression. Sports Medicine 55, 1449–1466. 10.1007/s40279-025-02204-7

Kawanishi, K., Nariyama, Y., Anegawa, K., Tsutsumi, M., Kudo, S., 2022. Changes in tibial nerve stiffness during ankle dorsiflexion according to in-vivo analysis with shear wave elastography. Medicine (United States) 101, E29840. 10.1097/MD.0000000000029840

Konrad, A., Gad, M., Tilp, M., 2015. Effect of PNF stretching training on the properties of human muscle and tendon structures. Scand J Med Sci Sports 25, 346–355. 10.1111/sms.12228

Konrad, A., Tilp, M., 2014. Increased range of motion after static stretching is not due to changes in muscle and tendon structures. Clinical Biomechanics 29, 636–642. 10.1016/j.clinbiomech.2014.04.013

Koo, T.K., Hug, F., 2015. Factors that influence muscle shear modulus during passive stretch. J Biomech 48, 3539–3542. 10.1016/j.jbiomech.2015.05.038

Kubo, K., Ishigaki, T., Ikebukuro, T., 2017. Effects of plyometric and isometric training on muscle and tendon stiffness in vivo. Physiol Rep 5, 1–13. 10.14814/phy2.13374

Le Sant, G., Nordez, A., Andrade, R., Hug, F., Freitas, S., Gross, R., 2017. Stiffness mapping of lower leg muscles during passive dorsiflexion. J Anat 230, 639–650. 10.1111/joa.12589

Miyamoto, N., Hirata, K., Miyamoto-Mikami, E., Yasuda, O., Kanehisa, H., 2018. Associations of passive muscle stiffness, muscle stretch tolerance, and muscle slack angle with range of motion: Individual and sex differences. Sci Rep 8. 10.1038/s41598-018-26574-3

Mukai, H., Umehara, J., Saeki, J., Yanase, K., Wang, Z., Tateuchi, H., Ichihashi, N., 2025. Tibial nerve stiffness is related to maximum angle of ankle dorsiflexion. Preprint. bioRxiv. 10.1101/2025.07.20.665743

Nakamura, M., Ikezoe, T., Kobayashi, T., Umegaki, H., Takeno, Y., Nishishita, S., Ichihashi, N., 2014. Acute effects of static stretching on muscle hardness of the medial gastrocnemius muscle belly in humans: An ultrasonic shear-wave elastography study. Ultrasound Med Biol 40, 1991–1997. 10.1016/j.ultrasmedbio.2014.03.024

Nakamura, M., Ikezoe, T., Umegaki, H., Kobayashi, T., Nishishita, S., Ichihashi, N., 2017. Changes in passive properties of the gastrocnemius muscle-tendon unit during a 4-week routine static-stretching program. J Sport Rehabil 26, 263–268. 10.1123/jsr.2015-0198

Rao, S.R., Saltzman, C.L., Wilken, J., Yak, H.J., 2006. Increased Passive Ankle Stiffness and Reduced Dorsiflexion Range of Motion in Individuals With Diabetes Mellitus. Foot Ankle Int 27, 617–622. 10.1177/107110070602700809

Riemann, B.L., DeMont, R.G., Ryu, K., Lephart, S.M., 2001. The Effects of Sex, Joint Angle, and the Gastrocnemius Muscle on Passive Ankle Joint Complex Stiffness. J Athl Train 36, 369–377.

Selcuk Can, T., Ozdemir, S., Yilmaz, B.K., 2022. Shear-Wave Elastography of Patellar Ligament and Achilles Tendon in Semiprofessional Athletes: Comparing With Nonexercising Individuals. Journal of Ultrasound in Medicine 41, 2237–2246. 10.1002/jum.15908

Takeuchi, K., Nakamura, M., Fukaya, T., Konrad, A., Mizuno, T., 2023. Acute and Long-Term Effects of Static Stretching on Muscle-Tendon Unit Stiffness: A Systematic Review and Meta-Analysis. J Sports Sci Med 3, 465–475. 10.52082/jssm.2023.465

Taniguchi, K., Shinohara, M., Nozaki, S., Katayose, M., 2015. Acute decrease in the stiffness of resting muscle belly due to static stretching. Scand J Med Sci Sports 25, 32–40. 10.1111/sms.12146

Weppler, C.H., Magnusson, S.P., 2010. Increasing muscle extensibility: A matter of increasing length or modifying sensation? Phys Ther 90, 438–449. 10.2522/ptj.20090012

